# Identifying Inheritance Patterns of Allelic Imbalance, using Integrative Modeling and Bayesian Inference

**DOI:** 10.64898/2026.03.28.714974

**Authors:** Stephanie H. Hoyt, Timothy E. Reddy, Raluca Gordân, Andrew S. Allen, William H. Majoros

## Abstract

Interpreting the effects of novel mutations on phenotypic traits remains challenging, particularly for *cis*-regulatory variants. For rare variants, individuals typically possess at most one affected copy of the causal allele, leading to allelic imbalance, and thus the ability to infer inheritance of allelic imbalance can inform genetic studies of phenotypic traits. While many methods for detection of allele-specific expression (ASE) exist, they largely focus on ASE in one individual. We show that performing joint inference across multiple individuals in a trio allows for simultaneously improving estimates of ASE and identifying its likely mode of inheritance. Our Bayesian approach has the benefit of being able to (1) aggregate information across individuals so as to improve statistical power, (2) estimate uncertainty in estimates, and (3) rank modes of inheritance by posterior probability. We demonstrate that this model is also applicable to other forms of imbalance such as allele-specific chromatin accessibility. Applying the model to ATAC-seq and RNA-seq from several trios, we uncover examples in which ASE can be linked to imbalance in chromatin state of *cis*-regulatory elements and to potential causal variants. As the cost of sequencing continues to decrease, we expect that powerful methodologies such as the one presented here will promote more routine collection of samples from related individuals and improve our understanding of genetic effects on gene regulation and their contribution to phenotypic traits.

## 1. Introduction

Interpretation of mutations in personal genomes remains challenging, particularly for *cis*-regulatory mutations, which have been strongly implicated in gene expression abnormalities [1–7]. A powerful approach to understanding the genetics of phenotypic traits is the study of co-segregation patterns of traits and genetic variants among related individuals, such as in a familial trio or larger pedigree. Indeed, co-segregation analyses have been found to substantially increase the ability to identify genetic causes of traits via identification of both inherited and *de novo* variants [8–11].

For rare variants affecting expression of genes, afflicted individuals often possess only one affected copy of the causal allele, leading to allelic imbalance, or *allele-specific expression* (ASE). The null expectation is that each copy of any given gene is expressed equally, such that in an RNA-seq sample, reads from the two alleles will be present in a roughly 1:1 ratio, but subject to noise due to sampling error. Many genes do not follow this pattern. One common form of this is *imprinting*, in which one parental haplotype is stably repressed in a non-random fashion, typically via heritable DNA methylation [12]. Another form is *random monoallelic expression*, in which either parental haplotype may be expressed, but on a per-cell basis, presumably due to epigenetic variation between individual cells [13]. ASE is particularly relevant for dosage-sensitive genes, which can mediate phenotypic effects for even subtle differences in expression [14].

Thus, for rare variants, the ability to identify patterns of co-segregation of trait status and allelic imbalance in relevant genes can be particularly informative. While gene expression aberrations can also be detected via between-individual comparisons, within-individual analyses via allelic imbalance have the virtue of being internally controlled for confounding factors such as genetic background and environmental effects, as the two gene copies being compared exist in the same nucleus and are subject to the same trans environment. In addition, ASE can often be detected with greater sensitivity than between-individual differences in gene expression, due to lower variance (PMID: 23300628). Moreover, ASE is generally indicative of *cis*-acting factors and thus can be relevant to the search for *cis*-regulatory causal variants [15–17].

Several methods currently exist for the detection of ASE. Many of these methods are formulated as *null hypothesis significance tests* (NHSTs) such as the binomial and beta-binomial tests, relying solely on the likelihood of the data at a single site under the null hypothesis (i.e., no ASE) to produce a p-value representing the statistical significance of any apparent imbalance under the assumption that imbalance is not truly present [18–22]. In contrast to NHST methods, Bayesian models can be used to directly assess evidence for both the null hypothesis (no ASE present) and alternate hypothesis (ASE present) within a single model, and due to their incorporation of prior probabilities, they facilitate direct estimation of *moderated effect sizes*--estimates that are automatically stabilized by model priors so as to mitigate false positives [23–25]. Bayesian methods can also provide full posterior probability distributions for effect sizes, enabling more thorough assessment of the evidence for ASE [26–31].

To date, existing methods have focused on assessing ASE in either isolated individuals [18, 26] or across extended populations [32–34], and have not extensively leveraged the shared information present in closely related individuals such as family members. While inheritance of ASE within a family and its co-segregation with traits of interest has been somewhat studied, research into the use of inheritance patterns of ASE in a formal statistical modeling framework has to date been minimal [34, 35]. When RNA samples are available from all individuals of a trio, a naive approach to identifying inheritance is to assume independence between individuals, classify each individual separately as to ASE status, and then to infer the inheritance pattern post hoc. As related individuals are in fact not independent, we expect that explicitly modeling the individuals and their expression data and shared genotypes may allow for more accurate inference of inheritance patterns and effect sizes, and ideally to better facilitate downstream efforts at identifying candidate causal mutations.

We thus propose a novel Bayesian model, *TrioBEASTIE* (*Trio-aware Bayesian Estimation of Allele-Specific Transcription by Integrating Evidence*), that extends our earlier model, BEASTIE [26], by jointly modeling allelic gene expression and per-gene-copy affected status across a mother, father, and offspring. Through detailed modeling of Mendelian transmission probabilities, *de novo* mutation rates, and possible recombination in the presence of latent proximal or distal causal factors, we can assign a posterior probability to possible modes of inheritance for ASE in a gene, as well as a posterior for the null case. Thus, we can report the best-supported inheritance mode and its posterior probability, as well as the sub-optimal modes, and we can also sum across modes to estimate a marginal probability for any individual in the trio having ASE. As the posterior probabilities are conditional on all the data in the trio, they effectively aggregate or “borrow strength” across multiple individuals in both assigning inheritance patterns and estimating effect sizes when multiple individuals are affected, leading to more accurate estimates than independence-based models.

Using a detailed, empirical trio-based simulator, we show that on simulated data the joint modeling approach outperforms an independence model in both identifying the true mode of inheritance and estimating the magnitude of ASE. We further illustrate the utility of this model by applying it to both ATAC-seq and RNA-seq data from two trios extensively characterized by the Thousand Genomes [36], Geuvadis [37], and CEPH [38] projects and demonstrate that imbalance in gene expression can be linked to imbalance in chromatin state of *cis*-regulatory elements, identifying putative causal variants within those elements.

## 2. Results

### 2.1 Overview of TrioBEASTIE

TrioBEASTIE is a probabilistic graphical model that estimates allele-specific activity in a gene (ASE) or open chromatin region (ASA) simultaneously across a trio, using read counts from heterozygous sites. TrioBEASTIE uses 11 modes of inheritance, for the 11 different ways we consider it possible for individuals in the trio to be affected (Fig. 1). These modes include a null (all unaffected) mode, modes where the child is the only affected individual, modes where the parent is affected and the child does or does not inherit, and modes where the parent is affected and recombines. We use modes that assume at most one affected copy in an individual and at most one affected parent. When evaluating each of these 11 modes of inheritance, the affected status of the individuals is fixed (Fig. 2 a-b, Methods 4.1, Supplementary Table 1). This means that when we sample values of θ (the effect size), this sampling is influenced only by the read counts of the individuals defined to be affected. This also means that TrioBEASTIE returns a posterior probability for each mode of inheritance (Fig. 2c). This allows a user to compare between modes and assess confidence in both classification and θ estimate. Information about additional parameters and priors can be found in Methods section 4.2 and Supplementary Methods 1.1.

**Figure 1:**
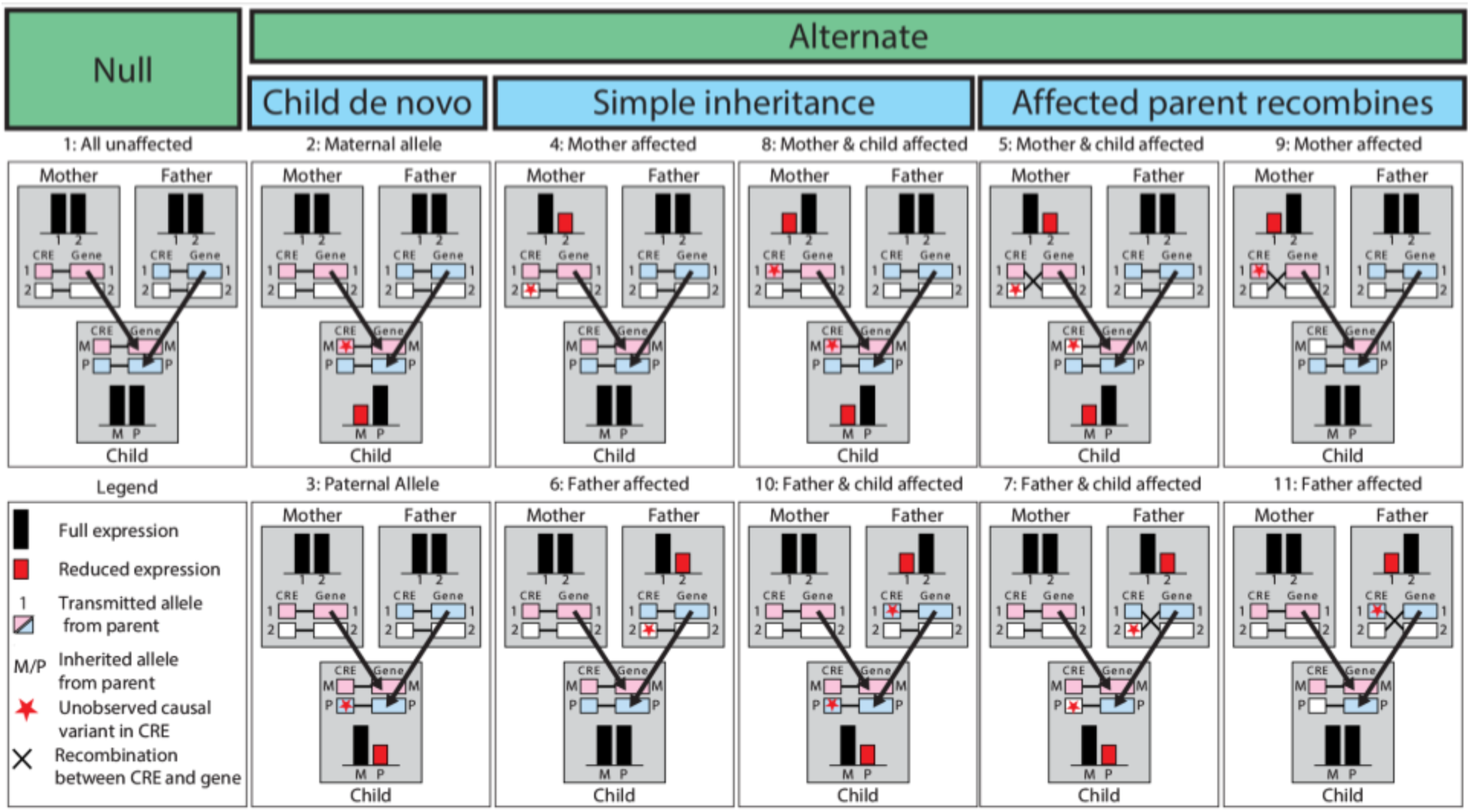
Visual descriptions of 11 modes of inheritance. The model detects ASE using gene expression read counts, but the ASE is being passed down through the inheritance of an unobserved causal element, likely in a cis-regulatory region (labeled red star in CRE). Red denotes an affected copy resulting in decreased expression (increased expression is also detectable but not illustrated); M=maternal, P=paternal. Modes 5,7,9, and 11 describe a recombination in the affected parent between the unobserved causal element and the gene body, such that the child inherits the gene allele associated with full expression, but the causal variant associated with reduced expression. Thus, the child has reduced expression on the opposite allele as the equally affected parent.

**Figure 2:**
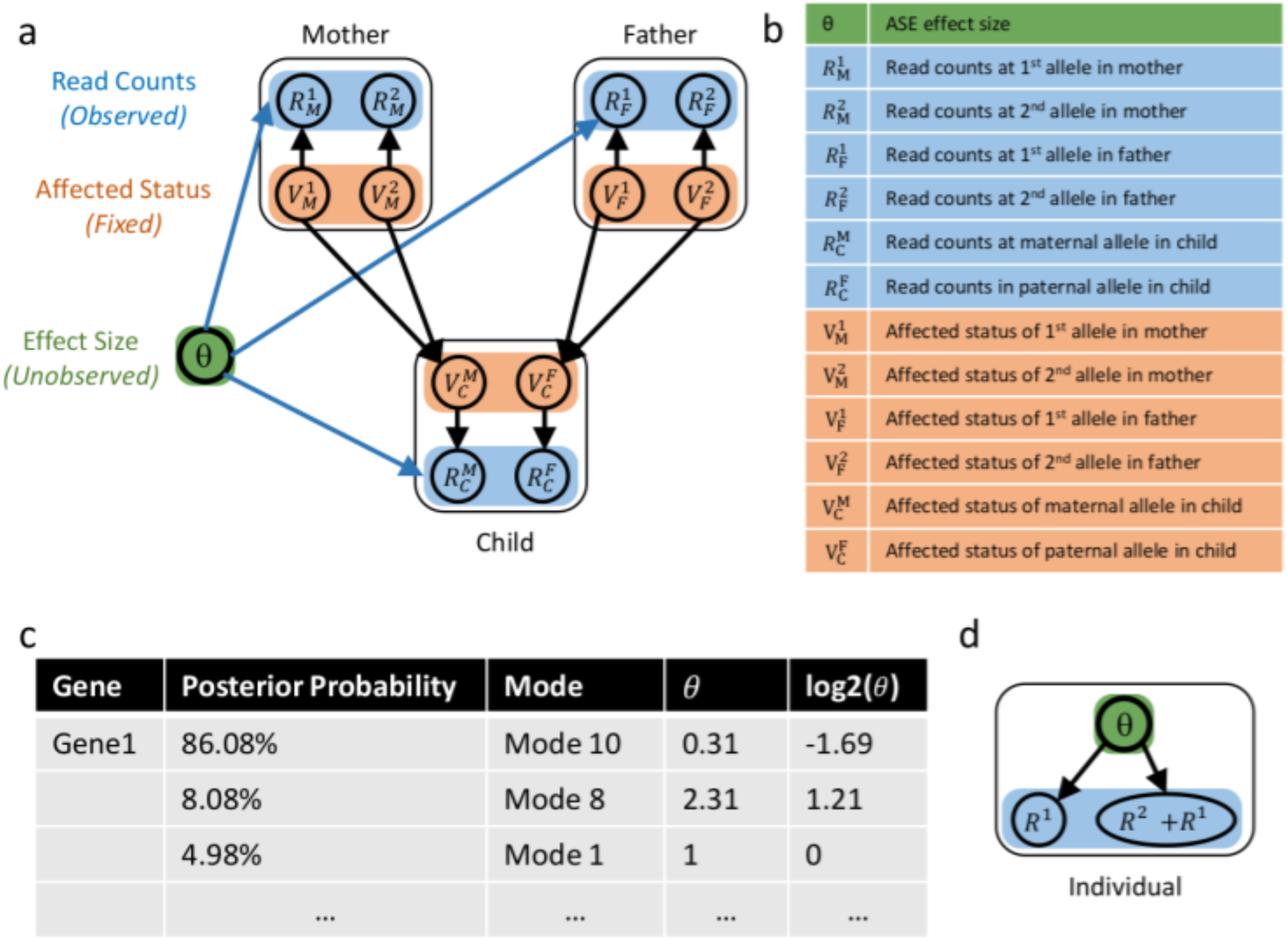
a) TrioBEASTIE is built up of 11 models (one for each mode of inheritance), which can be represented in a single graphical model by replacing fixed binary affected statuses with V variables. b) Model term definitions, including effect size, read counts, and affected status in each individual. c) For each gene, the output includes the normalized posteriors of each mode and the effect size. d) TrioBEASTIE can be compared to a baseline independence model, which estimates effect size based on read counts in a single individual.

### 2.2 TrioBEASTIE discriminates between simulated modes of inheritance

To assess TrioBEASTIE, we first generated simulated genes for each of our 11 modes of inheritance (Fig. 1). These genes are simulated by artificially ‘mating’ two individuals from the 1000 Genomes Project [36]: at each variable site, one allele is arbitrarily chosen to be passed down to the ‘child’. Genes are simulated over a range of values for the number of sites per gene, number of reads per site, and effect size of ASE (θ) (see Methods section 4.2). Allele-specific read counts are generated from a binomial distribution, where *p* = *θ* /(*θ* + 1).

TrioBEASTIE makes use of read count data to estimate the posterior probability of each mode of inheritance and sample θ values (Fig. 2 a, b). On this simulated data, TrioBEASTIE accurately detects ASE and distinguishes between modes of inheritance when formulated as a binary classification task of the correct inheritance pattern versus all other simulated patterns. All individual models achieve AUC greater than 0.7 when presented with sufficient data (Fig. 3 a-c, Supplementary Figure 1) (i.e., strong ASE and sufficient read depth across multiple sites). TrioBEASTIE has higher accuracy when θ is more extreme (meaning θ is farther from 1) and when there is more information (more reads per site and more sites per gene) (Fig. 3b, Supplementary Figure 1). With a more extreme θ, the model achieves higher accuracy with fewer reads (Fig. 3c). We assess performance per model only on genes where at least one individual defined to be affected in that model has at least one phased heterozygous site. Thus, we do not penalize models for inaccurate ASE calls when there are no supporting variants.

**Figure 3:**
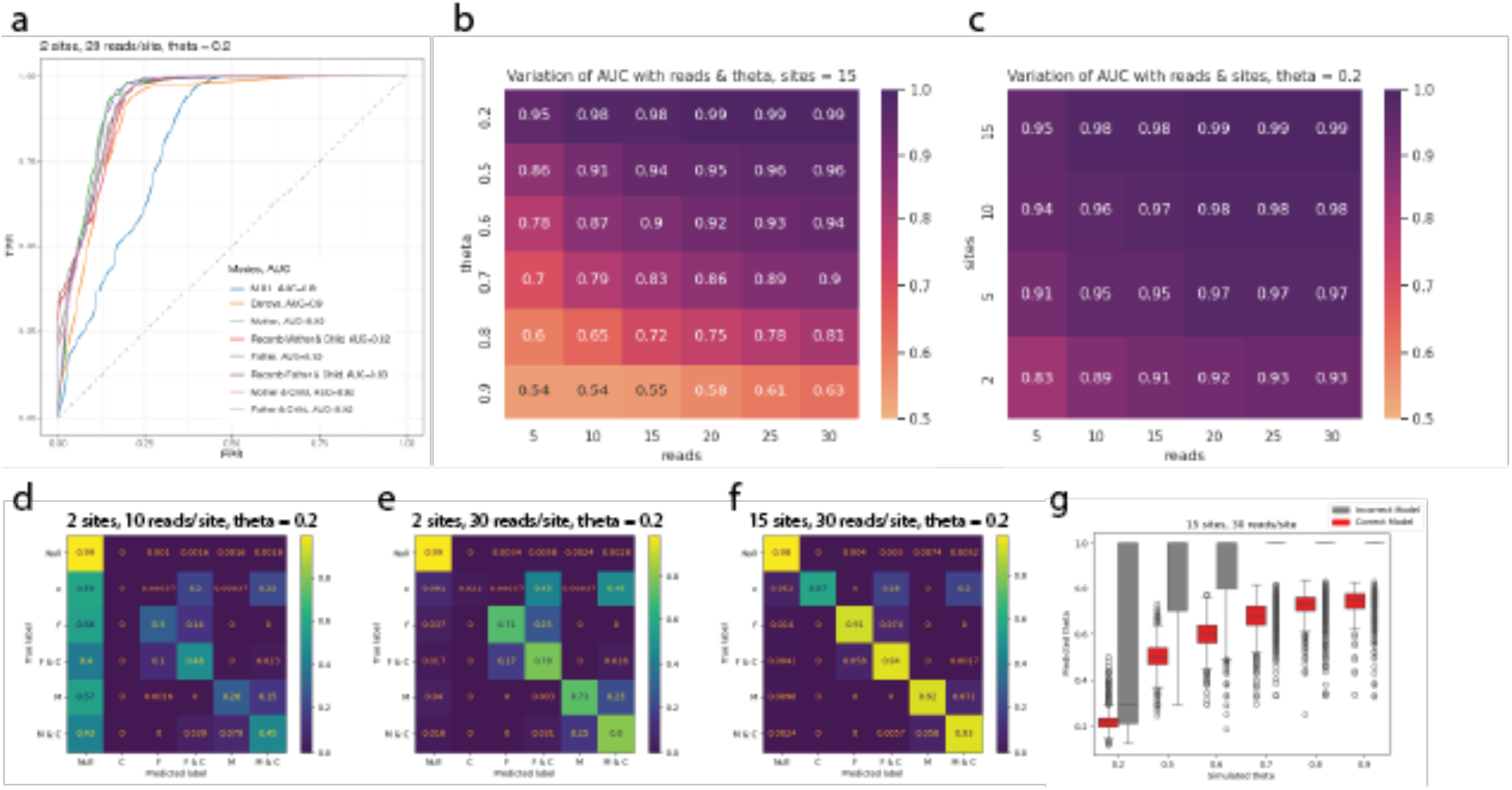
a) ROC curve for 11 modes of inheritance collapsed into 8 classes (*de novos* collapsed into one class, and parent only recombinations collapsed into modes where parents are affected and child does not inherit curve because they are indistinguishable). (b-c) Heatmaps of AUC values for model 10 (father affected and child inherits), summarizing binary classification results across some parameter combinations. (d-f) Confusion matrices showing that the main ‘confusion’ is to call null when we don’t have enough reads. Secondarily, we sometimes get confused about if just parent or parent + child are affected, but don’t get confused about if other parent is affected. ‘C’ row refers to both *de novo* modes, ‘F’ row is *Mode 6: Father affected*, ‘F &C’ row is *M10: Father and child affected*, ‘M’ row is *Mode 4: Mother affected*, and row ‘M & C’ row is *M8: Mother and child affected*. (g) Simulated vs predicted θ values in trio model, split by genes that were classified correctly (red) and genes that were classified incorrectly (gray).

### 2.2 TrioBEASTIE calls correct mode of inheritance and effect size θ per gene

In addition to assessing the performance of each model, we examined how often the correct mode of inheritance is identified for each simulated gene (Fig. 3 d-f, Supplementary Figure 2). This analysis focused on the relative probability of modes per gene, as opposed to the classification accuracy of a specific model. With sufficient read depth and number of sites, genes are most often called correctly (Fig. 3 e, f). Without sufficient information, the model is conservative; thus, genes are occasionally called as the null mode (Fig. 3d). This is by design and is due to the larger effect of shrinkage priors in the model when there is insufficient data to overcome the priors. Given sufficient information, the model can still exhibit confusion as to whether the parent and child or just the parent is affected, though at a low rate (Fig. 3e). The model does not show confusion as to which parent is affected, and all confusion decreases as we add more information (Fig. 3f). Some modes of inheritance are harder to discriminate, such as *de novos* where only the child is affected. These cases require additional information to call correctly a majority of the time (Fig. 3f).

In addition to assigning mode probabilities to each gene, TrioBEASTIE estimates a latent variable θ for the effect size of the ASE. Mode classification and identification of ASE are the primary goals of TrioBEASTIE, given that the specific effect size may not be as biologically or clinically relevant; it may be sufficient to know that an allele has reduced expression. Given this, we do still see that estimates of θ in simulated genes are generally close to the simulated true values, given sufficient information (Fig. 3g, Supplementary Figure 3). Estimated values are closer to the simulated truth when θ is extreme (farther from 1) and when there is more information. The most deviant θ estimates come from genes incorrectly classified into the null mode and thus assigned the point null (θ = 1). θ estimates are specific to the model that samples them, so we would not expect an estimate of θ to be accurate if the incorrect model is used.

### 2.3 TrioBEASTIE meets or exceeds baseline model accuracy

After determining TrioBEASTIE is accurate, we then endeavored to show it is more accurate than existing approaches which identify ASE in single individuals and combine calls across a trio post hoc. We compared TrioBEASTIE to a simple Bayesian model we call the independence model, which makes use of the trio phasing information, but only considers read counts from one individual (Fig. 2d). The independence model does not define the affected status of individual alleles the way TrioBEASTIE does; instead ASE calls are made by thresholding the posterior θ values. We also compare TrioBEASTIE to the binomial test, which returns a p-value at a single site. We first compare these models on their ability to call ASE in a single individual; we subset to genes where the child has at least one phased heterozygous site, so TrioBEASTIE and baseline models can be fairly compared. When estimating the presence of ASE in the child, TrioBEASTIE is as or more sensitive and specific than the independence model in 121 out of 144 tested parameter combinations (Fig. 4a-c). In the few cases where AUC is higher in the independence model, the difference is not significant by the Delong test [39, 40] (Fig. 4b).

**Figure 4:**
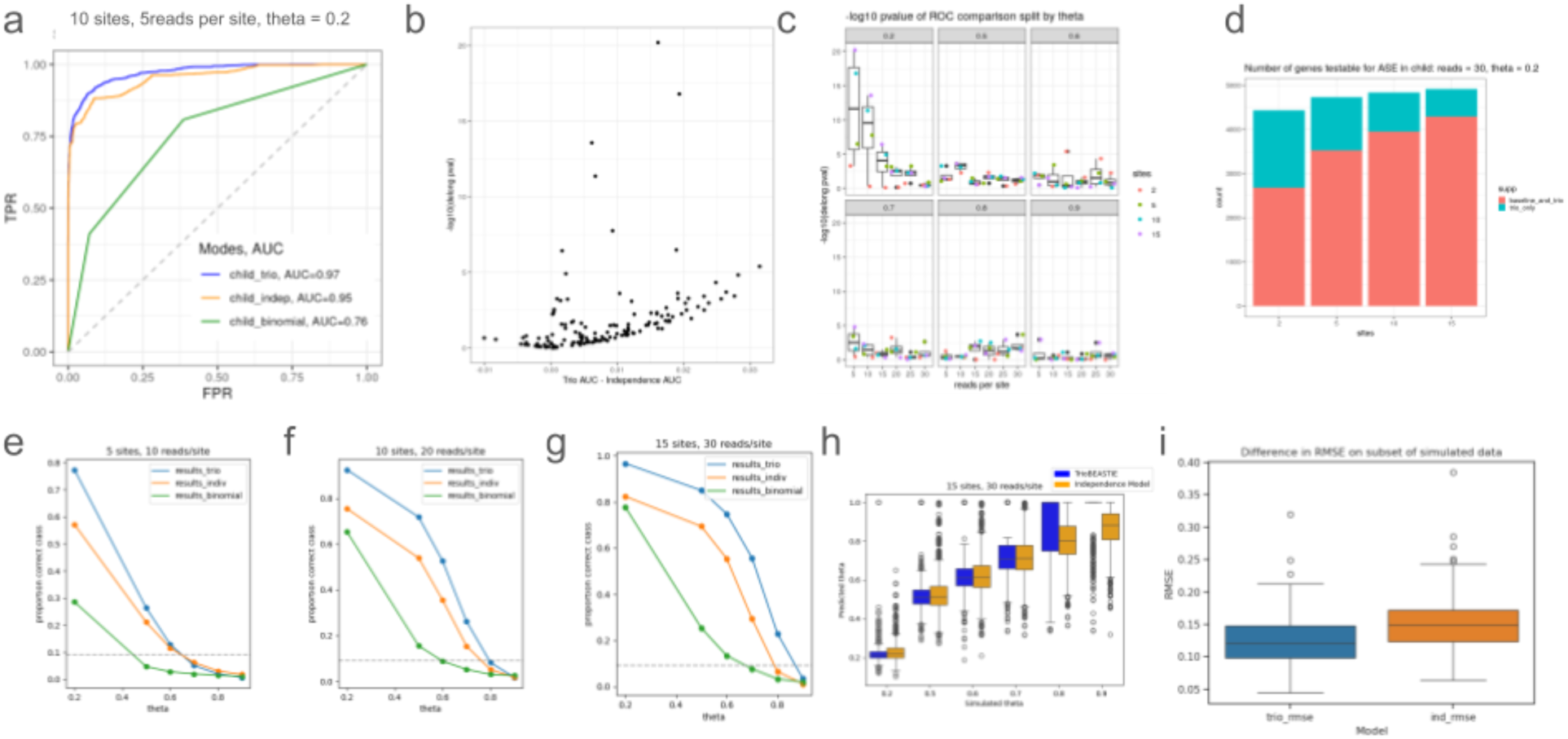
Always looking at ASE in the child, using genes simulated to be null or one of the simple modes of inheritance. (a) ROC curve comparing trio to baselines at a specific parameter combination where we see a difference between trio & independence mode. (b) Difference in AUC values between trio and independence model, and significance of those differences by the delong test for all parameter combinations. In few (7/144) cases where the independence model has higher AUC, the difference is non-significant. (c) Significance of AUC value differences split by all parameter combinations. Most significant differences are when θ is extreme (0.2) and read counts per site are small. (d) Stacked bar chart showing the number of genes trio vs baseline (independence, and binomial test) can run on. This shows the advantage of trio model because it can provide ASE estimates for more genes. It does this in an individual without direct genetic or RNA evidence by inferring ASE status from the status of related individuals. Baseline models only work on genes where there is direct evidence (ie at least 1 phased het in the individual of interest). (e-g) accuracy on reconstructing simulated mode of inheritance: 8-way classification accuracy for trio model and 6-way for baseline models (tested on ASE status per individual, instead of per copy). Trio is accurate if highest probability mode matched the truth.

TrioBEASTIE AUC values have the most significant difference from the independence model when θ is most extreme, and when read counts are low (Fig. 4c). This suggests that TrioBEASTIE is most useful when there is not enough information in the child to call ASE without additional data from a parent. Both TrioBEASTIE and the independence model perform better than the binomial test (Fig. 4a); the latter is limited by not combining information across sites and does not estimate the degree of the allelic imbalance (θ).

Using the trio model also allows us to assess the probability of ASE in the child, even when there are no phased heterozygous sites in the child, due to Mendelian transmission probabilities (Fig. 4d). This advantage in the additional number of genes for which TrioBEASTIE can assess probability of ASE is particularly clear when there are fewer sites in a gene, so that the chance of the child having reads at a phased heterozygous site is lower. Conditional on the child not having information, TrioBEASTIE has high accuracy in distinguishing null genes from those in which the child is affected, using parental information, but is reduced to random guessing when distinguishing between *parent only affected* and *parent affected, child inherits* modes (Supplemental Figure 4). This is expected, absent data in the child, as the Mendelian transmission probability of a child inheriting ASE from an affected parent is 0.5.

In addition to comparing accuracy in a single individual, we can also compare the models on their ability to call or reconstruct the correct mode of inheritance. For TrioBEASTIE, this is an 8-way classification task, requiring the most probable model to be the true model, considering the affected status of individual alleles (with the exception of the same mode collapsing done in ROC curves). For both baseline models this is a 6-way classification task, requiring the true affected individuals to be called correctly, but not considering the per-allele affected status of those individuals. TrioBEASTIE has a higher 8-way classification accuracy than either baseline (independence model or binomial test) in 6-way classification (Fig. 4 e-f). Classification accuracy for all models increases as the total amount of information increases. This shows how TrioBEASTIE shares information across related individuals to fit expected patterns of inheritance.

We can also compare effect size estimates in the child between TrioBEASTIE and the independence model; the binomial test does not generate an effect size. TrioBEASTIE θ estimates are comparable to those in the independence model, on genes in which the child is affected (Fig. 4g). In this analysis, because of the explicit classification performed by TrioBEASTIE, θ values estimated by the independence model are being compared to values from TrioBEASTIE which are either estimated during sampling or set to 1 in the case of a point null. This leads to an abundance of null θ values from TrioBEASTIE, particularly in cases with less information, where TrioBEASTIE makes greater use of null priors, as in Fig. 3h. In order to compare θ estimation between these models, without disadvantaging TrioBEASTIE, we considered a specialized subset of simulated data. In this subset, we consider genes from one mode of inheritance (mother affected and child inherits), we require both affected individuals to have an equal non-zero number of phased heterozygous sites, and we always use θ from the correct mode of inheritance, even if it is not the most likely. In addition to removing the barrier of classification for TrioBEASTIE, these criteria ensure we are looking at genes where the trio model has access to exactly twice as much information as the independence model, when the independence model is looking at the child. By subsetting to these data where TrioBEASTIE is designed to have an advantage, we can confirm if the model is working as expected and making use of all available information. We then compare the root-mean-squared-error (RMSE) of θ between the two models (Fig. 4i). As expected, the RMSE is significantly lower in TrioBEASTIE than in the independence model (two-sided Wilcoxon test p-value = 2.98e-25).

For independence model we threshold all ASE calls at P(ASE) >= 0.9; call anything >= 0.9 as having ASE and anything below as no ASE for each individual. Then build mode of inheritance out of three binary calls. If it matchs the simulated truth, then mark accurate. Similarly for binomial test, but thresholding p-value < 0.05. (h) Simulated vs predicted θ models comparing between trio model (purple) and independence model (green). We can see that the trio model is hindered by the discretization of θ that comes with the classification, but when it is actually estimating θ instead of assigning it to the point null, the results are comparable to the independence model. (i) RMSE difference between trio and independence model, on specialized subset of simulated data to show that trio model is able to take advantage of additional information that independence model does not have access to. two-sided Wilcoxon test p-value = 2.98e-25.

### 2.4 TrioBEASTIE identifies ASE genes in the CEPH1463 pedigree

To demonstrate the application of our model to future clinical data, we applied TrioBEASTIE to two genetically independent trios from the CEPH 1463 pedigree [38] (Fig. 5a). We refer to these as the NA12878 trio (including NA12878 and her parents, NA12891 and NA12892) and the NA12877 trio (including NA12877 and his parents, NA12889 and NA12890). These trios can be taken as examples of how our model can identify patterns of inheritance of ASE and allele-specific accessibility (ASA) and connect those calls to other genomic annotations that may lead to discovery of potential causal variants. For this analysis we use genotypes from the NYGC [41], and RNA-seq and ATAC-seq data from [42]. All preprocessing of sequencing data is done with the ENCODE pipelines [43] (see Methods 4.4).

**Figure 5:**
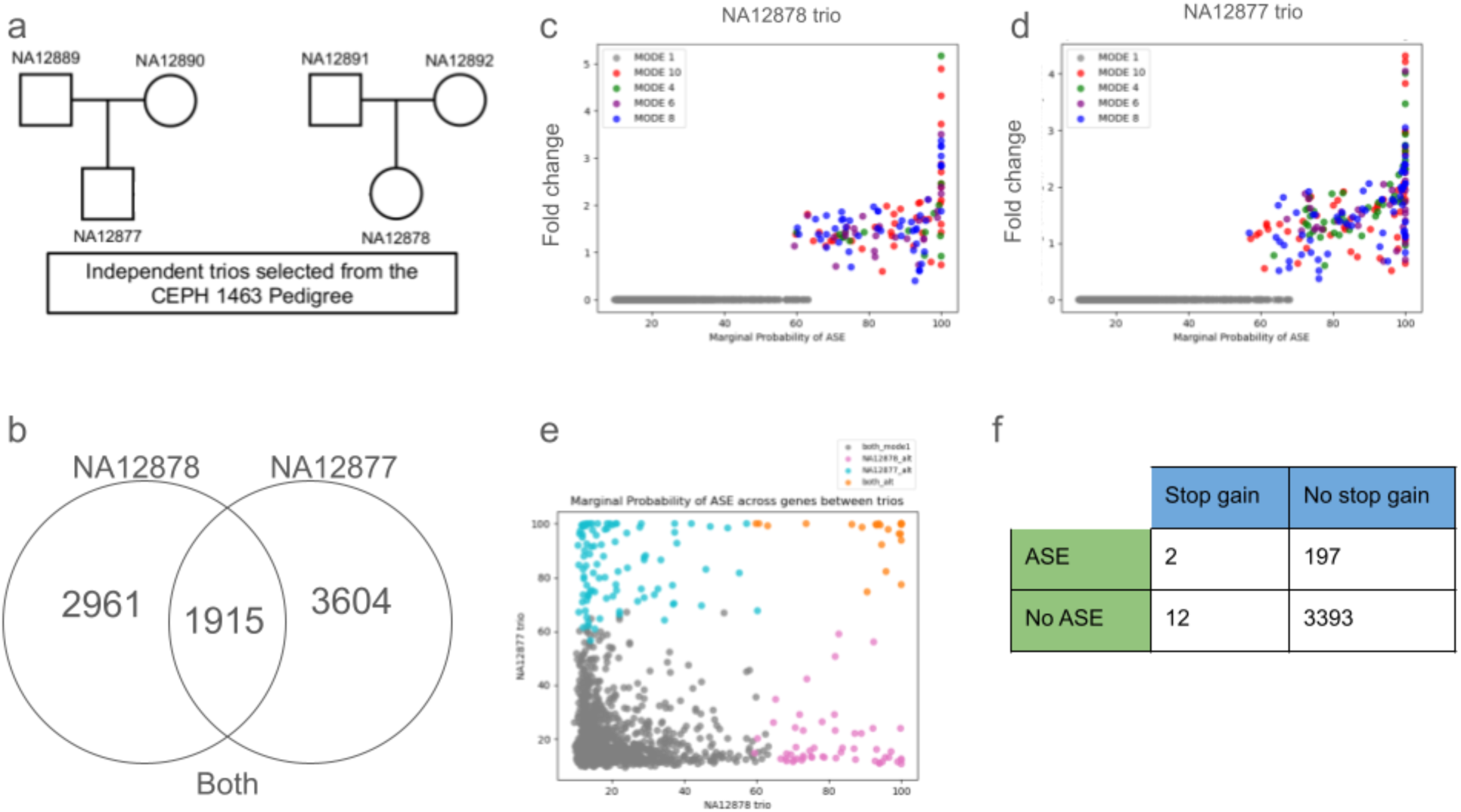
(a) Pedigree of trios. Ticks on bottom represent 11 children which are not used in this analysis. (b) Venn diagram showing number of genes testable for ASE in each trio (passing read depth threshold of at least 10 reads in at least 1 individual across all sites). Intersection is genes that go into panel E (c-d) Plots of genes w/ highest probability of ASE in NA12878 (c) and NA12877 (d) CEPH 1463 trios. (e) Comparison between trios. Genes which show ASE in both trios, may have a greater probability of underlying biological role (as opposed to random ASE), or could be due to shared variants between individuals. (f) LOF enrichment, defining LOF as a stop gain variant. Requires the variant to be in an individual predicted to be affected to go in ‘ASE & stop gain’ box.

There are 11,382 genes in the NA12878 trio that have at least one heterozygous site with overlapping RNA-seq reads in at least one individual in the trio, allowing us to run TrioBEASTIE. To remove underpowered genes (those predicted to be null due to lack of read depth information or signal), we apply a read depth filter requiring at least 10 reads in at least 1 individual across all sites. We do not apply a filter to any positive ASE genes because they are clearly powered to detect a signal. After applying this filter, we have 2,961 genes with enough data to reliably measure ASE (Fig. 5b). 132 of these are ASE genes, meaning the most likely mode is not the null *all unaffected* mode, and 22 genes have a 100% marginal probability of ASE. The ASE genes come entirely from *simple inheritance* modes, where one of the parents is affected, and the child either does or does not inherit the ASE (Fig. 5c). Approximately 4.46% (132/2961) of the genes we consider testable for ASE show high probability of ASE. This is in line with our empirical prior of 0.04 that an individual allele will be affected, leading to ASE (Methods 2.2). The greatest fold change values come from genes with 100% marginal probability of ASE. The top 5 genes with the highest fold changes in expression between the alleles in the NA12878 trio are IGKV3 (log2 fold change of most likely mode = 5.17), PABPC1 (4.89), RPS28P7 (4.32), RPSA2 (3.72), and SLFN5 (3.51). In all of these examples, the predicted affected individuals have read counts from only one allele.

In the NA12877 trio, we apply the model to 11,215 genes. After the same additional filtering steps were applied to the NA12878 trio, we have 3,604 genes with sufficient read depth to measure ASE (Fig. 5b). 200 of these are ASE genes, and 27 have a 100% marginal probability of ASE. All genes showing ASE are from *simple inheritance* modes (Fig. 5d). Some of these are ‘ambiguous genes’ where we can detect ASE but cannot determine the exact mode of inheritance due to a lack of information in some individuals. For example, in the NA12877 trio, there are no RNA-seq reads overlapping any phased heterozygous SNPs in the child within the gene body of SLFN5. We see strong ASE in the mother but cannot differentiate between the two modes: *mother affected, child doesn’t inherit,* and *mother affected, child inherits,* because we cannot measure ASE in the child. Thus, the marginal probability of at least 1 individual having ASE in this gene is 100%, and the marginal probability of the father having ASE is 100%, but the marginal probability of the child having ASE in this example is 49.64%. Each of the two modes of inheritance have their own effect sizes, which are approximately inverses of each other; the direction of θ is defined in terms of the ordering of the alleles, and we don’t know how to order the alleles if we don’t know if the child is affected. The genes in the NA12877 trio with the highest fold changes are PPIAP22 (log2 fold change 4.32), RPS28P7 (ambiguous, absolute value log2 fold changes for two modes are 4.22 and 4.06), CHI3L2 (4.06 and 19.36), HMGA1 (16.25 and 3.84), and RN7SL5P (3.84) (Fig. 5d). In this data, we do not see any genes for either trio for which the most likely mode of inheritance involves a *de novo* mutation or a recombination. This is not unexpected, given the rarity of *de novo* mutations and recombinations, in combination with the rarity of ASE.

To validate expected relationships between loss-of-function variants and the presence of ASE, we perform an enrichment test to see if ASE genes have more variants predicted to lead to a stop gain than non-ASE genes. In the NA12877 trio, we see that the stop gain variants are proportionally more prevalent in ASE genes (odds ratio = 2.87); however, the enrichment is not statistically significant (Fisher test p-value = 0.17), likely due to the rarity of the stop gain variants among genes tested for ASE (Fig. 5f). For the NA12878 trio, there are no stop gain variants in individuals predicted to show ASE (Fisher test p-value = 1) (Supplementary Table 2).

### 2.5 Some ASE genes are common to genetically independent trios

Given that these two trios are genetically independent, we do not expect them to share many ASE genes by chance. Therefore, for genes that do show ASE in both trios, we expect there to be some kind of biological function or shared variant explanation. There are 1,915 genes testable for ASE and passing the read depth filter, or exhibiting ASE, in both unrelated trios (Fig. 5b). We can compare the marginal probability of ASE across the trios to identify genes with recurrent allelic expression patterns. 28 of these genes are ASE genes in both trios, and 5 of those genes have a 100% marginal probability of ASE in both trios (orange points in Fig. 5e). These 5 genes are RPL29P4, CBX3, RPS28P7, RPS7P1, and SLFN5. Some genes show a high probability of ASE in only one of the trios. 54 genes have non-null modes of inheritance in the NA12878 trio and show no ASE (null mode) in the NA12877 trio (pink points in Fig. 5e). Similarly, 98 genes show non-null ASE in the NA12877 trio and null in the NA12878 trio (cyan points in Fig. 5e).

### 2.6 Allele-specific accessible peaks suggest causal variants for ASE

We applied TrioBEASTIE to ATAC-seq data generated from the same samples as the RNA-seq. The goal of this analysis is to identify ASA in regulatory regions, in a context where we can both see inheritance within the trio and provide a biological explanation or function for the ASA. In this analysis, we do not consider recombinations, which results in only 7 modes of inheritance. This is because, for ASE we expect recombinations to occur between the affected gene and an unobserved causal element. In an ASA analysis, we expect the causal element to be within the peak, so there is no room for a recombination. We use each peak in an IDR set as our genomic element, comparable to the canonical transcripts on which we evaluated ASE. In the NA12877 trio, there are 11,284 peaks testable with sufficient reads, and 839 are ASA peaks. (Fig. 6a). Out of 12,488 tested peaks passing read depth filtering in the NA12878 trio, 791 are ASA peaks (Supplemental Figure 5a).

**Figure 6:**
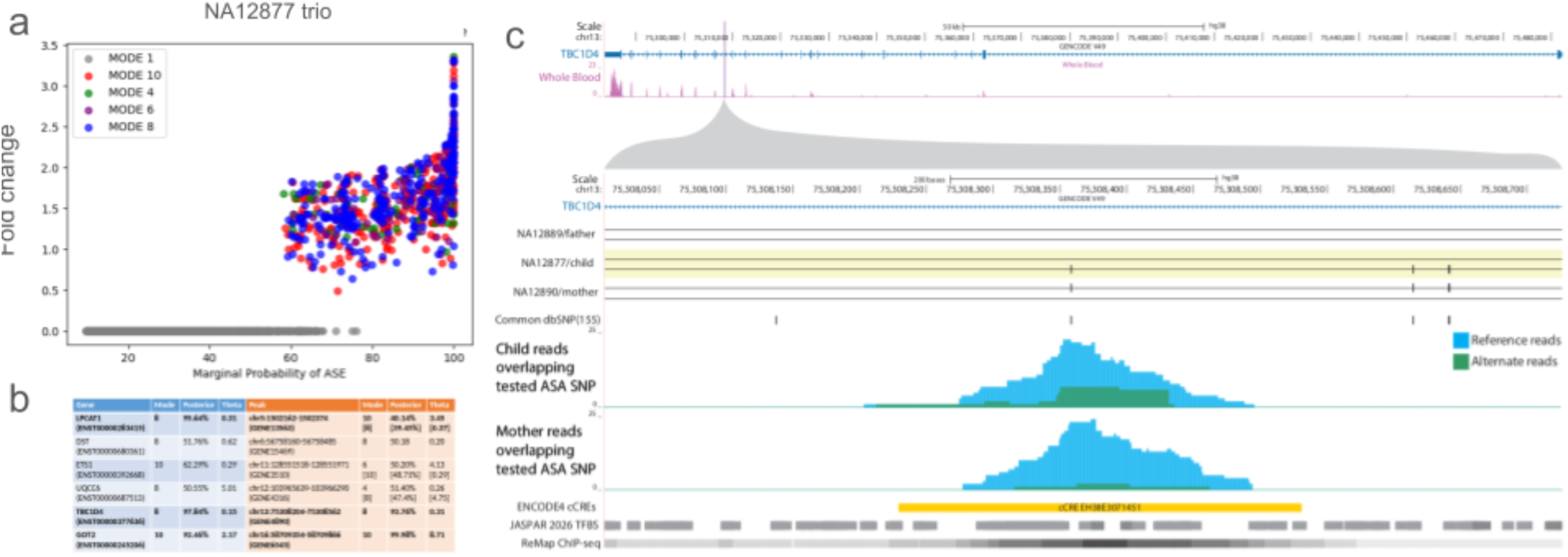
(a) Overview of ASA results in NA12877 trio (b) Concordance between ASE and ASA links in NA12877 trio (c) Example of ASE and ASA link at gene TBC1D4, which has an intronic ASA peak. The allele-specific reads are visible in blue and green overlapping an ENCODE cCRE. ASA is predicted in the mother and child, who share an alternate allele overlapping the ATAC-seq peak.

These ASA peaks are expected to have a downstream effect on gene expression. We identified potential regulatory networks by linking peaks to genes based on proximity. To prioritize ATAC-seq peaks more likely to have a regulatory role for a given gene, we focused on peaks either within the gene body or within 500 bp of the transcription start site. In the NA12877 trio, we were able to create 2,054 peak-gene links this way (across 1,397 genes and 1,972 peaks), of which 1,803 are cases where both peak and gene do not show ASE. Of the remaining 251 links where either the peak or gene does show ASE, 136 genes show ASE without corresponding ASA. Thus, in about half of cases of ASE with adjacent ATAC-seq peaks, the ASE can be explained by ASA in said peak. This number would be expected to rise if proximity requirements were loosened; this analysis prioritizes identifying peaks and genes very likely to be connected, as opposed to identifying all potential causal regulatory networks. There are 109 peaks showing ASA without downstream ASE, indicating that ASA at a single peak does not necessarily have a regulatory effect on adjacent genes. This could be due to the effect size of the ASE or due to the gene being regulated by multiple elements.

Of the greatest interest, there are 6 links where both the peak and gene show non-null ASE (Fig. 6b). If these links make up a true regulatory network where the ASA leads to ASE, we expect the same individuals to be affected in both contexts, and to show an effect in the same direction (i.e. decreased expression or accessibility on the same haplotype). Indeed, the mode and direction of θ are largely concordant between the regions; in 3/6 links the most likely mode for each matches, and in the other 3/6, the nearly equivalently likely ASA mode (shown in brackets) matches the most likely ASE mode. What may have originally appeared as a lack of concordance in those 3 links is likely due to low confidence in the most probable mode, due to insufficient read information. That these links are largely concordant in both mode and direction of effect size suggests that the ASA and ASE are inherited on the same haplotype and could be causally linked.

In the NA12878 trio, we have 1,601 peak-gene links (across 1,106 genes and 1,549 peaks). The majority (1,442/1,601) of these links are cases where both peak and gene do not show allele-specific activity, as expected. Out of the remaining 159 links where at least the peak or gene shows allele-specific activity, there are 73 and 82 cases where just the gene or just the peak shows allele-specific activity, respectively. There are 4 peak-gene links where both genomic regions show allele-specific activity (Supplemental Figure 5b).

Taking the link between TBC1D4 and an intronic peak in the NA12877 trio as an example, we can visualize the ASA and ASE within a genomic context using genome assembly hg38 in the UCSC Genome Browser [44] (Fig. 6c). The ASA peak overlaps ENCODE cCRE EH38E3071451 [45–48]. In separate data from the GM12878 cell line, this cCRE is classified as a transcription factor cCRE. There are intact Hi-C loops between the cCRE and TBC1D4. These factors all suggest a regulatory role for this ASA peak, which is also predicted by rE2G [49]. The peak overlaps many transcription factor binding sites predicted by JASPAR [50] with CREB binding being highly experimentally validated; however, the specific SNP tested in this peak does not overlap any binding sites. To propose specific variants that could be leading to the ASA, we consider those shared between the individuals predicted to be affected by TrioBEASTIE. There is one such site in this peak: chr13:75,308,359, known as rs1560540 [51] and heterozygous in the mother and child. This site is a known expression quantitative trait locus for TBC1D4 across many cell types [52]. When considering the ASE at this gene, we can see the signal is strong (Fig. 6b); however, there are no shared loss-of-function variants [53] that could explain the ASE. With the ASE unexplained by any variation within the gene body, this further supports the causal role of the ASA at the intronic peak.

## 3. Discussion

A key benefit of modern biotechnology in the post-genomic era is the ability to assay, rapidly and at relatively low cost, multiple aspects of human genetics and molecular physiology including genetic, epigenetic, and transcriptomic states, all of which have utility in promoting a better understanding of the genetics of human traits. Indeed, given the increased potential for data generation, an important recent trend within genomics broadly has been toward integrative modeling of multiple types and sources of data, in the hopes of exploiting synergistic relationships between them [54–56].

In this study, we showed that integrative modeling of genetic and transcriptomic data jointly across multiple related individuals can effectively exploit such synergies, resulting in substantial improvements in detecting gene expression abnormalities and their inheritance patterns across generations. This was made possible through detailed modeling of Mendelian transmission genetics together with Bayesian inference to produce robust estimates of effect sizes and provide uncertainty measures for those estimates. The key features of Bayesian modeling that come to bear in this setting are the ability to seamlessly integrate information from multiple observations in a principled, probabilistic way based on likelihoods; the ability to stabilize estimates of effect sizes by leveraging prior information through prior probability distributions; and the ability to derive posterior probabilities for individual modes of inheritance, effectively summarizing in a quantitative and directly interpretable way how well the data support each inheritance pattern.

We believe the ability to ascertain likely inheritance patterns of endophenotypes such as gene expression or epigenetic state has enormous potential in dissecting molecular mechanisms that can contribute to traits. However, while co-segregation of gene expression defects and trait status can be informative as to which genes may be contributing to a traits, such co-segregation will often not identify a definitive cause without additional evidence. Gene expression can be altered by a range of different genetic and epigenetic factors, including variants within a gene (e.g., loss-of-function mutations that trigger *nonsense-mediated decay*: NMD) as well as those potentially outside the affected gene (e.g., mutations in a transcriptional enhancer or promoter). For ASE estimation, variants within a gene are used merely to assign sequencing reads to gene copies and are not in general assumed to be causal for any imbalance in expression. Additional effort is needed to identify candidate causal mutations to explain the ASE, and while variant effect predictors can accurately assess some types of coding mutations (such as pre-termination codons triggering NMD), noncoding mutations can be much harder to definitively link to gene expression defects.

Allelic imbalance in chromatin has been suggested as an effective means of prioritizing variants in single individuals [57]. As demonstrated by our analyses of the two CEPH trios, our framework for modeling RNA-seq data in a trio is equally applicable to epigenetic data from assays such as ATAC-seq. In these example cases we detected possible co-segregation of ASE in genes and allele-specific chromatin accessibility (ASA) for cis-regulatory elements (CREs) that likely regulate those genes. Given the much smaller sizes of CREs and our current understanding of how accessibility is regulated via sequence determinants of nucleosome positioning [58, 59], variants within CREs exhibiting imbalance are much more likely to be causal for the imbalance than is the case for variants within ASE genes. In addition, many variants within known CREs have been or are in the process of being functionally annotated in public databases (e.g., https://data.igvf.org; https://catalog.igvf.org; [60]). As such, we posit that co-segregation of imbalance in multiple molecular traits, such as expression, accessibility, and transcription factor binding, will in general be more informative as to likely genetic causes than ASE alone.

While assays for epigenetic state (e.g., ATAC-seq, ChIP-seq) are not routinely performed in clinical settings at present, there are lower-throughput alternatives that can be used to validate small numbers of in silico predictions, such as luciferase assays for testing individual cis regulatory variants [61]. Given the relatively small number of ASE genes, we expect to observe co-segregating with the phenotype across a trio, these low-throughput options should be feasible. In addition, deep learning models for predicting allele-specific chromatin accessibility [62–64] while not expected to be as accurate as experimental measurements, may be useful for initial screening of variants for experimental validation.

While the modeling framework we have presented has clear utility, there are several areas for future improvement. First, while inheritance patterns across a trio can be informative for prioritizing genetic variants in trait studies, incorporating additional affected or unaffected family members can provide additional information [65, 66]. Given the flexibility of our Bayesian approach, we expect that accommodating larger pedigrees will be relatively straightforward, and indeed will enable more complete phasing of heterozygous sites in some cases [67]. Second, the model described here assumes that when multiple individuals exhibit ASE, the magnitude of that ASE is the same across individuals. While this assumption can easily be relaxed by introducing per-individual parameters for ASE magnitude, this incurs a risk of over-fitting.

However, that risk can likely be managed through hierarchical modeling and appropriate use of Bayesian shrinkage. Our model also assumes that each individual has at most one affected gene copy, and that at most one parent is affected; relaxing these assumptions can be accomplished by replacing our binomial likelihood with per-gene-copy negative binomial likelihoods for read counts. Modeling read counts directly in that way would also allow unrelated control samples to be included, which would in turn allow for testing simultaneously for either allelic imbalance or overall differences in expression of a gene as compared to controls.

Finally, we note that while genetically-driven ASE should follow the rules of Mendelian transmission genetics, it is known that cis regulatory variants can have low or incomplete penetrance due to multiple buffering mechanisms [67, 68], and when ASE contributes to complex traits via polygenic factors, the contribution of one gene to the phenotype can be very difficult to detect. Methods for incorporating ASE estimates and co-segregation patterns into burden tests and polygenic risk scores and similar approaches may thus be worthy of future investigation.

## 4. Methods

### 4.1 Model design

TrioBEASTIE is a probabilistic graphical model that represents ASE in a single gene across a trio (mother, father, and child), allowing inference across modes of inheritance of ASE (Fig. 1, Fig. 2a). A key innovation is the inclusion of the affected status for each individual and gene copy separately. The affected status is the presence or absence of ASE on a given haplotype. In this model, while the causal factor for ASE (whether genetic or epigenetic) might not be observed, and can be internal or external to the gene, the model can account for its presence via the latent affected status variables *V*^“^, for individual i and gene copy k (Fig. 2b). By jointly modeling affected status across all three individuals via a single joint distribution, the model can probabilistically infer the mode of inheritance of ASE by assigning posterior probabilities to each inheritance pattern.

Because an outgroup is not included, the represented modes of inheritance are limited to the cases in which dysregulation presents as allelic imbalance. Thus, we have eleven modes of inheritance (Fig. 1), consisting of one null mode where all individuals are unaffected, and ten alternate modes with at least one affected individual. We consider only cases with at most one affected parent and at most one affected copy per individual. We expect most genes showing ASE to fall into one of the *simple inheritance* modes, where a parent has an affected copy that is or is not transmitted to the child. Additionally, we model rarer events, including *de novo* ASE in the child, and recombinations in the parent between a putative (unobserved) causal element and the gene exhibiting ASE. Each of the eleven modes has a prior probability based on the total number of affected copies in the parents, and whether the mode requires a *de novo* event or recombination. In the case of *de novo* ASE, we see ASE in the child only. We model both directions of the ASE in the child (higher expression of the maternal or paternal haplotype).

We identify recombinations in a parent between the putative unobserved causal element and the gene body through read counts that are out of phase between the parent and the child, as follows. For example, in Mode 5 in Fig. 1, a recombination event has occurred between a distal cis-regulatory element and the gene body. As a result, the defective cis-regulatory allele regulates the non-transmitted gene copy in the mother and the inherited gene copy in the child. If the mother has biased read counts that are lower in the allele that the child does *not* inherit (identified through structural phasing), but the child also has biased read counts that are lower in the allele inherited from the mother, that suggests a recombination wherein the child inherited the correctly phased allele from the mother in the gene body, but inherited the other allele in the unobserved causal region, leading to ASE (Fig. 1).

We compare our model to two baseline models. The first is an *independence model* (Fig. 2d) that estimates ASE in each member of the trio separately, given a fixed phasing. We then run the model on all three individuals in the trio, separately, and assess the probability of ASE in each individual, from which we can make a binary call of the presence of ASE at a given probability threshold. Additionally, we compare to the binomial test, run at a single site within the gene in one individual at a time, using p = 0.5. We use the site with the deepest read depth; if there are multiple at equivalent read depths, we take the last one.

### 4.2 Posterior inference of effect size and mode of inheritance

TrioBEASTIE is implemented in Stan and run using RStan [69, 70] which performs Hamiltonian Monte Carlo sampling [71] using the No-U-Turn sampler [72]. The model is implemented as a collection of 11 models (Fig. 2a), one for each mode of inheritance, which allows an effect size to be estimated separately for each mode. In each, the *V*^“^ variables are fixed according to the defined affected statuses of each individual. These variables are what differentiate the models and modes of inheritance (Supplementary Table 1). We have two observed variables for each individual: the read counts of each allele *R*^“^ (Fig. 2b). We quantify ASE using a θ value, which represents the odds between the read counts on the two alleles in an individual, aka the effect size. We use a single θ value across all affected individuals, thus assuming that when an imbalance is present in multiple individuals, it has the same magnitude. We estimate θ for a given mode of inheritance, meaning when running the model for mode 8, θ would be influenced by the read counts at the two alleles in the mother and the child, but not by the read counts in the father. For the null model, we fix θ to a point null (θ = 1) and evaluate the likelihood of the reads at that point. The output (Fig. 2c) can be interpreted by looking at the posterior probability and θ of each mode of inheritance individually or can be summarized by calculating the marginal probability of ASE. The marginal probability of ASE is the sum of the posterior probabilities across all modes where at least one individual is affected.

### 4.3 Realistic simulation of read count data from 1000 Genomes Project individuals

We generate simulated data to test our model by artificially ‘mating’ two individuals from the 1000 Genomes project (HG00097, HG00096) [36]. For a given site in the human genome, we take the genotypes at that site for those two individuals. We then, without loss of generality and simplifying the representation, always assume each parent passes its maternal gene copy to the child. We skip triple homozygous sites. We then generate read counts for these individuals based on the simulated mode of inheritance, θ, and the number of reads per site. We simulate the number of sites per gene by repeating this process until we reach the desired number before moving on to the next gene. We simulate data for all modes of inheritance and varying θ, read depths, and reads per site so we can assess our model across these parameters.

Models are assessed on simulated data using various classification accuracy tasks. For each model and each parameter combination, we generate an ROC curve (analysis done in R version 4.4.0). This tasks the models with differentiating the genes simulated to match that model’s mode of inheritance from all other genes (binary classification). For a gene to be included in the ROC curve, there must be evidence in the simulated gene, meaning at least one phased heterozygous site in at least one affected individual, where here ‘affected’ means one of the individuals defined to be affected for the given mode of inheritance. For this analysis, we collapse the *de novo* modes into each other because they are mathematically indistinguishable, except for the direction of θ. Additionally, we collapse each of the ‘parent affected and recombines, child does not inherit’ into its corresponding ‘parent affected, child does not inherit’ mode, because the recombinations are mathematically not detectable without the child being affected, so those modes are indistinguishable. When generating confusion matrices (analysis done in Python version 3.13.2) of classification accuracy per class of simulated genes (as opposed to per model), we again collapse the *de novo* modes but do not include any recombination modes. By ‘collapsing two modes’, we mean that we consider any genes simulated to be from either of the two modes to be in the same class, and to be classified correctly if they are placed into either group.

When comparing TrioBEASTIE to the baseline models, we again make ROC curves, however, these are not per mode of inheritance model. The task is to classify the ASE status in the child correctly. Given this, we only consider simulated genes where the child has at least one phased heterozygous site so that there is information accessible to the baseline models. In TrioBEASTIE, we test if the most likely model is any one where the child is predicted to be affected. For each of the baseline models, we test if the point estimate of ASE reaches a significance threshold in the child. For the independence model, this is a probability of ASE greater than or equal to 0.9, and for the binomial test, this is a p-value of less than 0.05. In order to assess the difference between TrioBEASTIE and the independence method across parameters, we perform a two-sided Delong test on the AUC values [39, 40].

In addition to binary classification, we assess TrioBEASTIE and baseline models on 8-way and 6-way classification tasks, respectively. For each simulated gene, we assess TrioBEASTIE on whether the most likely model matches the true simulated mode of inheritance, similar to the analysis shown in confusion matrices. This is an 8-way classification, as there are 11 modes, but we collapse modes the same as in the ROC curve analysis, leaving us with 8.

Additionally, in this analysis we consider whether or not a gene is ambiguous, meaning there is sufficient information in the parents but the child or in the child but not the parents. In these genes, the posterior probability is split between two modes because there is not sufficient information. We call these correct if the most probable model is either of the ambiguous modes. This requires not only getting the affected status of each individual correct, but also the affected status of the individual alleles. In the baseline models, we do not have allele-level information, so we instead perform an easier 6-way classification just requiring the significance level of each affected individual to meet the classification threshold.

When comparing RMSE between TrioBEASTIE and the independence model, we focus on a subset of the simulated data to make the comparison fairer. We consider only genes where the mother and child are both affected; in order for the trio model to have an advantage, more than one individual needs to be affected. We also require both the mother and child to have an equal non-zero number of phased heterozygous sites; the difference between the models will be more visible when the trio model has exactly twice as much information at the independence model. We use the TrioBEASTIE θ estimate from the correct model, even if it isn’t the most likely; the independence model does not have to do an additional classification task, so we take that barrier away from the trio model.

### 4.4 Data acquisition and preprocessing

We use the first and second generations of the CEPH 1463 pedigree [38] and treat these six individuals as two independent, unrelated trios. We refer to these as the NA12878 trio (including NA12878 and her parents, NA12891 and NA12892) and the NA12877 trio (including NA12877 and his parents, NA12889 and NA12890). Genotypes are from the NYGC [41] (https://ftp.1000genomes.ebi.ac.uk/vol1/ftp/data_collections/1000G_2504_high_coverage/working/20220422_3202_phased_SNV_INDEL_SV/). RNA-seq and ATAC-seq are from [42]. Using bcftools [73], we concatenate and filter the NYGC vcf files into a vcf file per sample that contains only heterozygous biallelic SNPs across all autosomal chromosomes. We generate trio-level vcf files that include all sites where at least one individual in the trio is heterozygous. We are only interested in regions with heterozygosity because it is required to distinguish between alleles and measure allelic activity.

To analyze the RNA-seq, we run a version of the ENCODE RNA-seq pipeline that runs STAR with WASP instead of just STAR [43, 74] (https://github.com/s-hoyt/rna-seq-pipeline-wasp) in order to filter out reads with mapping bias. After running the pipeline, we run Picard to mark duplicates, using the random scoring method based on previously published best practices [75]. We then use samtools mpileup at all of the sites in our trio vcf file to count the reference and alternate reads overlapping each heterozygous site. We ignore any sites where all three individuals in a trio are heterozygous, as these are not phasable. We run TrioBEASTIE per autosomal gene, defining a gene as anything in Ensembl hg38 whose feature type is a gene and is on chromosomes 1 through 22 [76] (https://hpc.nih.gov/refdb/dbview.php?id=1094). We only keep sites from our trio vcf files that are overlapped by at least 1 read after running samtools mpileup and that lie within an exon of the canonical transcript of the gene, as defined in Ensembl hg38. In order not to not double count reads that span multiple sites, we thin sites such that we do not keep any within a read length (200bp) of each other. We exclude any sites that are not phasable by our trio phaser (Supplementary Methods 1.2). We additionally exclude any genes that fall within the ENCODE blacklist exclusion filter [46–48] (downloaded on October 9, 2024 from https://www.encodeproject.org/files/ENCFF356LFX/) and the UCSC unusual regions [44]; any gene counts or specific gene rankings listed in the results are from after the filtering.

ATAC-seq reads are processed using the ENCODE ATAC-seq pipeline and WASP [43, 77] (https://github.com/ENCODE-DCC/atac-seq-pipeline); the ENCODE alignment-only pipeline is used for each alignment step within the WASP framework. After the final bam file of reads passing WASP is generated, the full ENCODE pipeline is used to generate peaks. Using IDR optimal narrow peaks from each sample, we generate a trio peak set by taking the union of peaks across the trio and then merging any peaks that overlap by more than 100 base pairs. Bam files for each biological replicate are merged, and samtools mpileup is used at all of the sites in our trio vcf file to count the reference and alternate reads. Blacklist exclusion filtering is included as a step of the ENCODE pipeline, and we add a filter for the UCSC unusual regions.

### 4.5 Variant Effect Prediction analysis

For each trio, we consider the potential impact of variants overlapping genes tested for ASE. We use Ensembl Variant Effect Predicter (VEP) [53] to identify stop gain variants, which we believe are the most directly connected to ASE. We run VEP on all variants heterozygous in at least one individual in the trio overlapping a canonical exon of a tested transcript. This includes variants not tested for ASE for any of the following reasons: triple het site, no overlapping reads, or too close to another site. To link the annotations from VEP with the ASE results, we first merge gene-level ASE results with site information so each variant is linked to the transcript it overlaps, regardless of whether that site was actually used in ASE evaluation. Then, those sites are merged with VEP annotations by position only (ignoring the gene that the VEP annotation is associated with). We keep all genes tested for ASE even if they have no VEP annotations and subsequently consider those genes as having no stop gain variants. All genes are subjected to the same read filtering as in other analyses: either the most likely mode must be non-null, or there must be 10 reads in at least one individual for us to be confident that the gene is not null simply due to lack of power. We test for enrichment of stop gain variants by making a Fisher table of gene counts, broken into presence/absence of ASE and presence/absence of stop gain variants. The stop gain variant must be in the individual predicted to be affected for it to count as an ASE+stop gain case, otherwise it counts as no ASE+stop gain. We perform a one-sided Fisher test, looking for the ASE genes to have significantly more stop gain variants.

## Availability

https://github.com/s-hoyt/trioBEASTIE

## Author contributions

Model development and analysis: SHH, WHM Conceptualization and supervision of the work: WHM, ASA, RG

## Acknowledgements and Funding

Research reported in this publication was supported in part by the National Institute of General Medical Sciences of the National Institutes of Health under award number 1R35-GM150404 to W.H.M., and by NIH under award number RM1-HG011123 to T.E.R. and A.S.A. and R.G. Content is solely the responsibility of the authors.

## References

1. Lee, P.H., et al., Principles and methods of in-silico prioritization of non-coding regulatory variants. Hum Genet, 2018. 137(1): p. 15–30.

2. Zhang, G., et al., DiseaseEnhancer: a resource of human disease-associated enhancer catalog. Nucleic Acids Res, 2018. 46(D1): p. D78–D84.

3. Li, X., et al., The impact of rare variation on gene expression across tissues. Nature, 2017. 550(7675): p. 239–243.

4. Zhang, F. and J.R. Lupski, Non-coding genetic variants in human disease. Hum Mol Genet, 2015. 24(R1): p. R102–10.

5. Spielmann, M. and S. Mundlos, Looking beyond the genes: the role of non-coding variants in human disease. Hum Mol Genet, 2016. 25(R2): p. R157–R165.

6. Castel, S.E., et al., Modified penetrance of coding variants by cis-regulatory variation contributes to disease risk. Nat Genet, 2018. 50(9): p. 1327–1334.

7. Corradin, O. and P.C. Scacheri, Enhancer variants: evaluating functions in common disease. Genome Med, 2014. 6(10): p. 85.

8. Malmgren, H., et al., Diagnostic yield of 1000 trio analyses with exome and genome sequencing in a clinical setting. Front Genet, 2025. 16: p. 1580879.

9. Gupta, P., et al., Familial co-segregation and the emerging role of long-read sequencing to re-classify variants of uncertain significance in inherited retinal diseases. NPJ Genom Med, 2023. 8(1): p. 20.

10. Jarvik, G.P. and B.L. Browning, Consideration of Cosegregation in the Pathogenicity Classification of Genomic Variants. Am J Hum Genet, 2016. 98(6): p. 1077–1081.

11. Hansen, R.D., A.F. Christensen, and J. Olesen, Family studies to find rare high risk variants in migraine. J Headache Pain, 2017. 18(1): p. 32.

12. Bartolomei, M.S., Genomic imprinting: employing and avoiding epigenetic processes. Genes Dev, 2009. 23(18): p. 2124–33.

13. Kravitz, S.N., et al., Random allelic expression in the adult human body. Cell Rep, 2023. 42(1): p. 111945.

14. Veitia, R.A., J. Zschocke, and J.A. Birchler, Gene Dosage Sensitivity and Human Genetic Diseases. J Inherit Metab Dis, 2025. 48(4): p. e70058.

15. Campino, S., et al., Validating discovered Cis-acting regulatory genetic variants: application of an allele specific expression approach to HapMap populations. PLoS One, 2008. 3(12): p. e4105.

16. Renganaath, K., et al., Systematic identification of cis-regulatory variants that cause gene expression differences in a yeast cross. Elife, 2020. 9.

17. Shih, C.H. and J. Fay, Cis-regulatory variants affect gene expression dynamics in yeast. Elife, 2021. 10.

18. Rozowsky, J., et al., AlleleSeq: analysis of allele-specific expression and binding in a network framework. Mol Syst Biol, 2011. 7: p. 522.

19. Glinos, D.A., et al., Transcriptome variation in human tissues revealed by long-read sequencing. Nature, 2022. 608(7922): p. 353–359.

20. Hasenbein, T.P., et al., Allele-specific genomics decodes gene targets and mechanisms of the non-coding genome. Nucleic Acids Res, 2025. 53(19).

21. Raghupathy, N., et al., Hierarchical analysis of RNA-seq reads improves the accuracy of allele-specific expression. Bioinformatics, 2018. 34(13): p. 2177–2184.

22. Castel, S.E., et al., Rare variant phasing and haplotypic expression from RNA sequencing with phASER. Nat Commun, 2016. 7: p. 12817.

23. Smyth, G.K., Linear models and empirical bayes methods for assessing differential expression in microarray experiments. Stat Appl Genet Mol Biol, 2004. 3: p. Article3.

24. Jaffrezic, F., et al., A structural mixed model for variances in differential gene expression studies. Genet Res, 2007. 89(1): p. 19–25.

25. Marot, G., et al., Moderated effect size and P-value combinations for microarray meta-analyses. Bioinformatics, 2009. 25(20): p. 2692–9.

26. Zou, X., et al., Bayesian estimation of allele-specific expression in the presence of phasing uncertainty. Bioinformatics, 2025. 41(6).

27. Skelly, D.A., et al., A powerful and flexible statistical framework for testing hypotheses of allele-specific gene expression from RNA-seq data. Genome Res, 2011. 21(10): p. 1728–37.

28. Nariai, N., et al., A Bayesian approach for estimating allele-specific expression from RNA-Seq data with diploid genomes. BMC Genomics, 2016. 17 Suppl 1(Suppl 1): p. 2.

29. Leon-Novelo, L.G., et al., A flexible Bayesian method for detecting allelic imbalance in RNA-seq data. BMC Genomics, 2014. 15(1): p. 920.

30. Pirinen, M., et al., Assessing allele-specific expression across multiple tissues from RNA-seq read data. Bioinformatics, 2015. 31(15): p. 2497–504.

31. Knowles, D.A., et al., Allele-specific expression reveals interactions between genetic variation and environment. Nat Methods, 2017. 14(7): p. 699–702.

32. Castel, S.E., et al., A vast resource of allelic expression data spanning human tissues. Genome Biol, 2020. 21(1): p. 234.

33. Fan, J., et al., ASEP: Gene-based detection of allele-specific expression across individuals in a population by RNA sequencing. PLoS Genet, 2020. 16(5): p. e1008786.

34. Chen, J., et al., A uniform survey of allele-specific binding and expression over 1000-Genomes-Project individuals. Nat Commun, 2016. 7: p. 11101.

35. McDaniell, R., et al., Heritable individual-specific and allele-specific chromatin signatures in humans. Science, 2010. 328(5975): p. 235–9.

36. Genomes Project, C., et al., A global reference for human genetic variation. Nature, 2015. 526(7571): p. 68–74.

37. Lappalainen, T., et al., Transcriptome and genome sequencing uncovers functional variation in humans. Nature, 2013. 501(7468): p. 506–11.

38. Dausset, J., et al., Centre d’etude du polymorphisme humain (CEPH): collaborative genetic mapping of the human genome. Genomics, 1990. 6(3): p. 575–7.

39. DeLong, E.R., D.M. DeLong, and D.L. Clarke-Pearson, Comparing the areas under two or more correlated receiver operating characteristic curves: a nonparametric approach. Biometrics, 1988. 44(3): p. 837–45.

40. Sun, X. and W. Xu, Fast Implementation of DeLong’s Algorithm for Comparing the Areas Under Correlated Receiver Operating Characteristic Curves. IEEE Signal Processing Letters, 2014. 21(11).

41. Byrska-Bishop, M., et al., High-coverage whole-genome sequencing of the expanded 1000 Genomes Project cohort including 602 trios. Cell, 2022. 185(18): p. 3426–3440 e19.

42. Johnston, A.D., et al., Functional genetic variants can mediate their regulatory effects through alteration of transcription factor binding. Nat Commun, 2019. 10(1): p. 3472.

43. Hitz, B.C., et al., The ENCODE Uniform Analysis Pipelines. bioRxiv, 2023.

44. Casper, J., et al., The UCSC Genome Browser database: 2026 update. Nucleic Acids Res, 2026. 54(D1): p. D1331–D1335.

45. Moore, J.E., et al., An expanded registry of candidate cis-regulatory elements. Nature, 2026.

46. Kagda, M.S., et al., Data navigation on the ENCODE portal. Nat Commun, 2025. 16(1): p. 9592.

47. Consortium, E.P., An integrated encyclopedia of DNA elements in the human genome. Nature, 2012. 489(7414): p. 57–74.

48. Luo, Y., et al., New developments on the Encyclopedia of DNA Elements (ENCODE) data portal. Nucleic Acids Res, 2020. 48(D1): p. D882–D889.

49. Gschwind, A.R., et al., An encyclopedia of enhancer-gene regulatory interactions in the human genome. bioRxiv, 2023.

50. Ovek Baydar, D., et al., JASPAR 2026: expansion of transcription factor binding profiles and integration of deep learning models. Nucleic Acids Res, 2026. 54(D1): p. D184–D193.

51. Sherry, S.T., et al., dbSNP: the NCBI database of genetic variation. Nucleic Acids Res, 2001. 29(1): p. 308–11.

52. Consortium, G.T., The Genotype-Tissue Expression (GTEx) project. Nat Genet, 2013. 45(6): p. 580–5.

53. McLaren, W., et al., The Ensembl Variant Effect Predictor. Genome Biol, 2016. 17(1): p. 122.

54. Yang, J., et al., A Scalable Bayesian Method for Integrating Functional Information in Genome-wide Association Studies. Am J Hum Genet, 2017. 101(3): p. 404–416.

55. Sigurdsson, A.I., et al., Deep integrative models for large-scale human genomics. Nucleic Acids Res, 2023. 51(12): p. e67.

56. Baiao, A.R., et al., A technical review of multi-omics data integration methods: from classical statistical to deep generative approaches. Brief Bioinform, 2025. 26(4).

57. Morilla, L.M.O., et al., Leveraging allelic imbalance in accessible chromatin to prioritize putative causal variants. Trends Genet, 2026.

58. Struhl, K. and E. Segal, Determinants of nucleosome positioning. Nat Struct Mol Biol, 2013. 20(3): p. 267–73.

59. Chereji, R.V. and D.J. Clark, Major Determinants of Nucleosome Positioning. Biophys J, 2018. 114(10): p. 2279–2289.

60. Li, D., et al., The IGVF catalog-from genetic variation to function. Nucleic Acids Res, 2026. 54(D1): p. D1437–D1445.

61. Littleton, S.H. and S.F.A. Grant, Protocol to study cis-regulatory activity of GWAS loci for specific gene promoters in human primary astrocytes using luciferase reporter assay. STAR Protoc, 2024. 5(4): p. 103338.

62. Pampari, A., et al., ChromBPNet: bias factorized, base-resolution deep learning models of chromatin accessibility reveal cis-regulatory sequence syntax, transcription factor footprints and regulatory variants. bioRxiv, 2025.

63. Avsec, Z., et al., Advancing regulatory variant effect prediction with AlphaGenome. Nature, 2026. 649(8099): p. 1206–1218.

64. Zhang, Z., et al., Developing a general AI model for integrating diverse genomic modalities and comprehensive genomic knowledge. bioRxiv, 2025.

65. Cukier, H.N., et al., Exome sequencing of extended families with autism reveals genes shared across neurodevelopmental and neuropsychiatric disorders. Mol Autism, 2014. 5(1): p. 1.

66. Ganesh, S., et al., Whole exome sequencing in dense families suggests genetic pleiotropy amongst Mendelian and complex neuropsychiatric syndromes. Sci Rep, 2022. 12(1): p. 21128.

67. Blackburn, A.N., et al., Genotype phasing in pedigrees using whole-genome sequence data. Eur J Hum Genet, 2020. 28(6): p. 790–803.

68. Payne, J.L. and A. Wagner, Mechanisms of mutational robustness in transcriptional regulation. Front Genet, 2015. 6: p. 322.

69. Team, S.D., RStan: the R interface to Stan. 2025.

70. Team, S.D., Stan Reference Manual. 2026.

71. Betancourt, M.J. and M. Girolami, Hamiltonian Monte Carlo for Hierarchical Models. arxiv, 2013.

72. Hoffman, M.D. and A. Gelman, The No-U-Turn Sampler: Adaptively Setting Path Lengths in Hamiltonian Monte Carlo. Journal of Machine Learning Research, 2014.

73. Danecek, P., et al., Twelve years of SAMtools and BCFtools. Gigascience, 2021. 10(2).

74. Asiimwe, R. and D. Alexander, STAR+WASP reduces reference bias in the allele-specific mapping of RNA-seq reads. bioRxiv, 2024.

75. Castel, S.E., et al., Tools and best practices for data processing in allelic expression analysis. Genome Biol, 2015. 16(1): p. 195.

76. Dyer, S.C., et al., Ensembl 2025. Nucleic Acids Res, 2025. 53(D1): p. D948–D957.

77. van de Geijn, B., et al., WASP: allele-specific software for robust molecular quantitative trait locus discovery. Nat Methods, 2015. 12(11): p. 1061–3.

78. Milo, R. and R. Phillips, Cell biology by the numbers. 2015: CRC Press.

79. Acuna-Hidalgo, R., J.A. Veltman, and A. Hoischen, New insights into the generation and role of de novo mutations in health and disease. Genome Biol, 2016. 17(1): p. 241.

