## Supplementary text and figures for "Identifying Inheritance Patterns of Allelic Imbalance, using Integrative Modeling and Bayesian Inference"

### Methods:

#### 1.1 Additional model details

We consider only 11 modes of inheritance in this model, because to consider additional modes requires an outgroup or control expression value to compare each allele to. Instead, here we look only for imbalance between the two alleles. Additional modes that could be included in a future model are those where an individual has two affected copies (observed as total decrease in expression, or decrease in both copies), and cases where both parents are affected because that could lead to two affected copies in the child. The 11 modes of inheritance are defined using binary affected status variables (Supplementary Table 1).

| | $V_M$ | $V_F$ | $V_C$ | $P(V_C \mid V_M, V_F)$ | |
|---|---|---|---|---|---|
| 1 | 00 | 00 | 00 | $1-P_{denovo}$ | No affected alleles |
| 2 | 00 | 00 | 10 | $0.5P_{denovo}$ | De novo in child |
| 3 | 00 | 00 | 01 | $0.5P_{denovo}$ | De novo in child |
| 4 | 01 | 00 | 00 | $1-P_{recomb}$ | Mother affected, child does not inherit |
| 5 | 01 | 00 | 10 | $P_{recomb}$ | Mother affected and recombines, child inherits |
| 6 | 00 | 01 | 00 | $1-P_{recomb}$ | Father affected, child does not inherit |
| 7 | 00 | 01 | 01 | $P_{recomb}$ | Father affected and recombines, child inherits |
| 8 | 10 | 00 | 10 | $1-P_{recomb}$ | Mother affected, child inherits |
| 9 | 10 | 00 | 00 | $P_{recomb}$ | Mother affected and recombines, child doesn't inherit |
| 10 | 00 | 10 | 01 | $1-P_{recomb}$ | Father affected, child inherits |
| 11 | 00 | 10 | 00 | $P_{recomb}$ | Father affected and recombines, child doesn't inherit |

**Supplementary Table 1:** For each mode, we encode the affected status for each copy in each individual as 0 or 1. In the child, the first allele represents the maternal allele, and the second allele is the paternal allele. In the parents, the first allele is the allele passed on to the child, and the second allele is the non-inherited allele. In the parents, we arbitrarily define the maternal allele as the one passed on to the child and the paternal allele as the non-inherited copy. In the event of a recombination, the child inherits the paternal (second) allele instead.

Within the Stan model, if an individual is defined to be affected for a given mode, then we use p defined by the sampled theta within the binomial likelihood. Else, we use 0.5 as the expected proportion of reads from each allele as we are testing the mode where this individual does not have ASE. We allow $\log\theta$ to be greater than or less than zero, representing an increase or a decrease in expression relative to the unaffected allele, respectively.

We use the default 4 chains per model run to check for convergence using the R-hat statistic (convergence threshold set at 1.05). We generate 5000 samples per model run, of which half (2500) are warmup. θ and posteriors are both calculated using the posterior mean over all non-warmup samples. Within the model, we have fixed priors on recombinations (0.001), *de novo* mutations (0.0001), and on an individual allele being affected (0.04). Priors on recombination and *de novo* are based on published rates [78, 79]. The prior of an allele being affected is derived from previously observed rates of ASE across the CEU population within the

1000 Genomes Project [26]. Using the observed individual affected rate of 0.0768, we can write that 0.0768 = 2p(1-p) because we expect an individual to be heterozygous for an affected allele to exhibit ASE. We can then solve for p to get the rate of affected alleles, p = 0.04, which becomes the prior in our model.

We define θ = p/(1-p), where p is the parameter of the binomial test representing the proportion of reads that come from one parental gene copy versus the other. Our null expectation is that there will not be any difference between the read counts, ie, there will be no ASE, so we place a shrinkage prior on θ: $\theta \sim log_2 N(0,1)$. We calculate the likelihood of our observed data under a given mode of inheritance using the binomial likelihood summed across all three individuals: $P(R_M, R_F, R_C | V_M, V_F, V_C, \theta, \Phi) = \Sigma_i R^1{}_i \sim binom(R^1{}_i | p, R^1{}_i + R^2{}_i)$, where $p = \theta/(\theta+1)$ and $i \in \{M, F, C\}$. Read counts are per site in the gene, and θ is estimated by summing likelihoods across all sites. The posterior is: $P(R_M, R_F, R_C, V_M, V_F, V_C, \theta \mid \Phi) = P(R_M, R_F, R_C | V_M, V_F, V_C, \theta, \Phi) P(V_C | V_M, V_F) P(V_M) P(V_F) P(\theta)$ We then run the model for the remaining ten different possible alternate modes of inheritance, estimate ten different θs, and then calculate the posterior probability of all modes (null and alternate) using Bayes theorem: by dividing the likelihood of a given mode by the sum of the likelihoods of all modes.

$$P(V_M, V_F, V_C | R_M, R_F, R_C, \theta, \Phi) = \frac{P(R_M, R_F, R_C, V_M, V_F, V_C, \theta \mid \Phi)}{\Sigma_{V_M V_F V_C} P(R_M, R_F, R_C, V_M, V_F, V_C, \theta \mid \Phi)}$$

After running the model for all eleven modes, we estimate the posterior probability of each mode using Bayes' theorem. For any given MCMC sample, this looks like equation (2). Since we have likelihoods from 1000 MCMC samples, we generate a posterior by averaging across all of these samples, shown in equation (3).

$$\begin{aligned} & P(V_M^*, V_F^*, V_C^* | \boldsymbol{R_M}, \boldsymbol{R_F}, \boldsymbol{R_C}, \theta, \boldsymbol{\Phi}) = \\ & \frac{P(\boldsymbol{R_M}, \boldsymbol{R_F}, \boldsymbol{R_C}, V_M^*, V_F^*, V_C^*, \theta | \boldsymbol{\Phi})}{\Sigma_{V_M^*, V_F^*, V_C^*} P(\boldsymbol{R_M}, \boldsymbol{R_F}, \boldsymbol{R_C}, V_M^*, V_F^*, V_C^*, \theta | \boldsymbol{\Phi})} = \\ & \frac{P(\boldsymbol{R_M}, \boldsymbol{R_F}, \boldsymbol{R_C} | V_M^*, V_F^*, V_C^*, \theta, \boldsymbol{\Phi}) P(V_C^* | V_M^*, V_F^*) P(V_M^*) P(V_F^*) P(\theta)}{\Sigma_{V_M^*, V_F^*, V_C^*} P(\boldsymbol{R_M}, \boldsymbol{R_F}, \boldsymbol{R_C} | V_M^*, V_F^*, V_C^*, \theta, \boldsymbol{\Phi}) P(V_C^* | V_M^*, V_F^*) P(V_M^*) P(V_F^*) P(\theta)} = \\ & \frac{P(\boldsymbol{R_M}, \boldsymbol{R_F}, \boldsymbol{R_C} | V_M^*, V_F^*, V_C^*, \theta, \boldsymbol{\Phi}) P(V_C^* | V_M^*, V_F^*) P(V_M^*) P(V_F^*)}{\Sigma_{V_M^*, V_F^*, V_C^*} P(\boldsymbol{R_M}, \boldsymbol{R_F}, \boldsymbol{R_C} | V_M^*, V_F^*, V_C^*, \theta, \boldsymbol{\Phi}) P(V_C^* | V_M^*, V_F^*) P(V_M^*) P(V_F^*)} \end{aligned} \tag{2}$$

The posterior probability of mode $j$ for $j \in 1, 11$, calculated for each sample $i$ for $i \in 1, 1000$ and then averaged over all per-sample posteriors:

$$\begin{aligned} & \text{MCMC Posterior}(P(V_M^*, V_F^*, V_C^* | \boldsymbol{R_M}, \boldsymbol{R_F}, \boldsymbol{R_C}, \theta, \boldsymbol{\Phi})) = \\ & \frac{\Sigma_{i=1}^{1000} [P(V_M^*, V_F^*, V_C^* | \boldsymbol{R_M}, \boldsymbol{R_F}, \boldsymbol{R_C}, \theta, \boldsymbol{\Phi})]_i}{1000} = \\ & \frac{\Sigma_{i=1}^{1000} [\frac{P(\boldsymbol{R_M}, \boldsymbol{R_F}, \boldsymbol{R_C}, V_M^*, V_F^*, V_C^*, \theta | \boldsymbol{\Phi})}{\Sigma_{V_M^*, V_F^*, V_C^*} P(\boldsymbol{R_M}, \boldsymbol{R_F}, \boldsymbol{R_C}, V_M^*, V_F^*, V_C^*, \theta | \boldsymbol{\Phi})}]_i}{1000} = \\ & \frac{\Sigma_{i=1}^{1000} [\frac{P(\boldsymbol{R_M}, \boldsymbol{R_F}, \boldsymbol{R_C} | V_M^*, V_F^*, V_C^*, \theta, \boldsymbol{\Phi}) P(V_C^* | V_M^*, V_F^*) P(V_M^*) P(V_F^*) P(\theta)}{\Sigma_{V_M^*, V_F^*, V_C^*} P(\boldsymbol{R_M}, \boldsymbol{R_F}, \boldsymbol{R_C} | V_M^*, V_F^*, V_C^*, \theta, \boldsymbol{\Phi}) P(V_C^* | V_M^*, V_F^*) P(V_M^*) P(V_F^*) P(\theta)}]_i}{1000} \end{aligned} \tag{3}$$

When running the baseline independence model, we use CmdStan [70] with 5000 samples, of which 300 are warmup. A single θ estimate is calculated from the posterior median. The posterior probability of ASE is calculated by splitting the posterior distribution of θ into right and left tails at 1.2 and 1/1.2, respectively, and taking the maximum of the areas under the right and left tails. The baseline binomial test does not require any sampling.

1.2 Trio phaser

We phase genotypes per site between the three individuals in the familial trio. Simply put, we check from which parent the child could have inherited each of it's alleles. Phasing is only applied when at least one individual is heterozygous. This structural phasing method assumes there are no recombination events within the gene, there are no de novo mutations within the gene, and there are no sequencing errors (after stringent quality filtering). Meaning, if we observe any sites that violate these assumptions, they are not phased and are ignored in any subsequent analysis. Additionally, triple heterozygous sites can not be structurally phased and are ignored. After structural phasing, we can simply sum over all sites in a gene

### Results:

2.1 TrioBEASTIE performance on simulated data

Here we provide additional plots showing model performance across parameter combinations. Supplemental figure 1a supports the conclusions of Figure 3 a-b which says that more read depth information increases AUC, but that theta signal has a larger effect. This is also seen in the additional ROC curves in Supp Fig. 1b-c where AUC decreases as theta moves towards the null, even with a lot of reads.

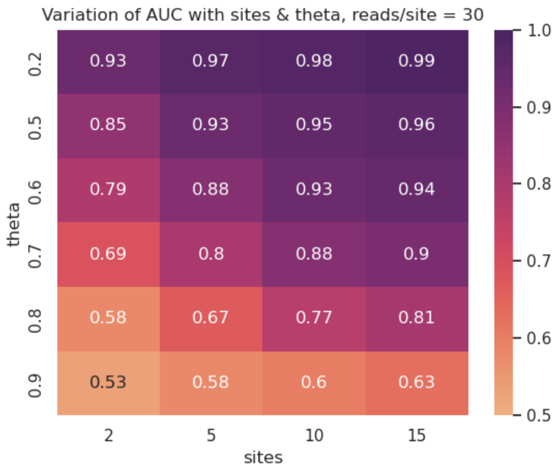


**Supplementary Figure 1:** Heatmap of AUC values for model 'father affected and child inherits', with reads/site fixed at 30 to observe effect of theta and number of sites on AUC.

In addition to assessing TrioBEASTIE classification per model, we also asses accuracy per gene, visualized through confusion matrices. Here, we show extended confusion matrices covering all 11 modes. In these plots, no modes are collapsed. We can see that *de novo* genes are frequently misclassified, which is expected due to the priors. Recombination modes where the child is unaffected are always misclassified, because they are mathematically indistinguishable from the case where the parent does not recombine.

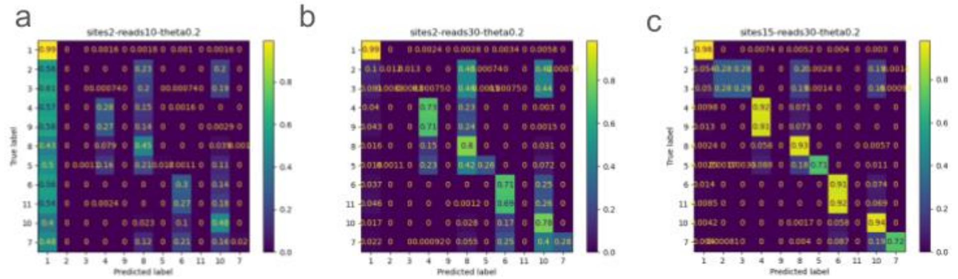


**Supplementary Figure 2:** Extended confusion matrices for the same parameter combinations shown in Figure 3. Sorted so that related modes are next to each (ie mother only affected is mode 4 and is next to mode 9, mother only affected and she has a recombination).

Beyond classification, TrioBEASTIE also estimates an effect size, which we can compare to the simulated effect size. This is shown for one combination of sites and reads per site in the main text. Here we provide results for additional parameter combinations.

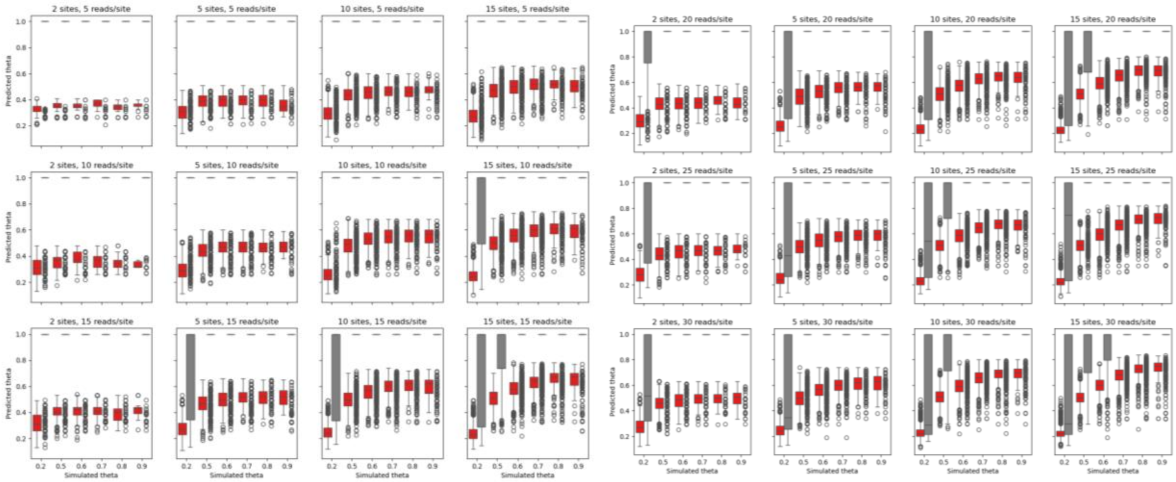


**Supplementary Figure 3**: Extended parameter combinations for theta expected vs observed estimates in simulated data. Needs a key: gray boxes are genes classified incorrectly (tends to be classification to the null mode with theta = 1) and red is classified correctly. We don't expect theta to be accurate if the mode is wrong. We can see sensitivity increase as we add more information, in the lower limit of θ's being estimated. When we have only 2 sites with 30 reads θ is never below 0.6 even when it is simulated to be 0.9. As we add more information, the range of θ estimates of more null genes get closer to true θ.

When extending TrioBEASTIE to individuals with no direct evidence, we see different accuracy based on the classification task. Using genes where there is no evidence in the child, but there is evidence in the parent we can either attempt to differentiate cases where the child is affected from null genes, or we can differentiate between cases where the parent is always affected but the child does or does not inherit the ASE. In the case of differentiating from null, the model has high accuracy (Supplemental Figure 4a). This is because if the parent is affected, there is a 50% chance the child inherits, which is much higher than a null case where the parent is unaffected so we can differentiate those genes. When the parent is always affected, the classification accuracy in the child drops to 50% because we now randomly guessing and we have no additional information beyond Mendelian inheritance patterns.

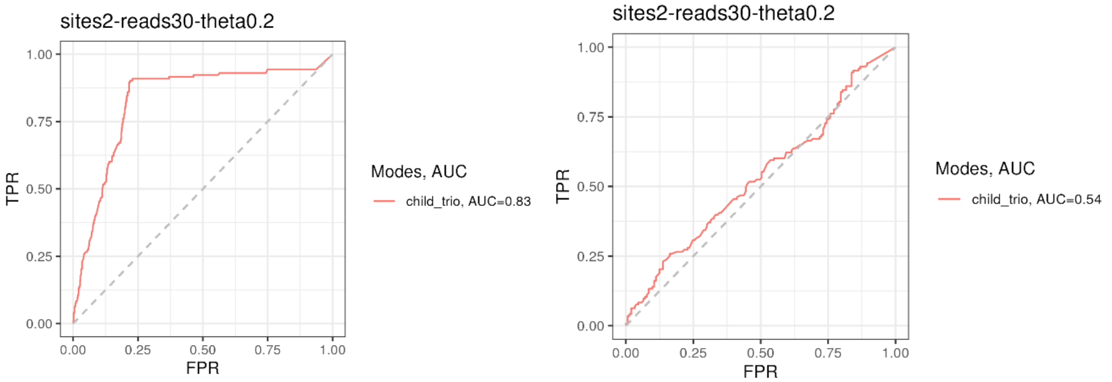


**Supplemental Figure 4: a)** ROC Showing trio model performance on trio only genes, when comparing true parent and child affected (cases) to null all unaffected + parent only affected (controls). **b)** ROC When comparing true parent and child affected (cases) to parent only affected (controls)

2.2 TrioBEASTIE identifies ASE and ASA in CEPH 1463 data

Figure 5 include stop gain variant enrichment for the NA12877 trio. Supplementary Table 2 shows the results for the corresponding analysis in the NA12878 trio. There are no stop gain variants in any ASE genes, for the individuals predicted to be affected. For this one-sided fisher test, the odds ratio is 0 and the p-value is 1.

**Supplementary Table 2:**

| NA12878 | Stop gain | No stop gain |
|---|---|---|
| ASE | 0 | 131 |
| No ASE | 15 | 2815 |

In figure 6, we focus on the NA12877 trio. Here are the corresponding results for the NA12878 trio. We identify similar number of ASA peaks (Supplemental Figure 5a). We are able to create 4 peak-gene links where both elements show allele-specific activity; however, only one of these links shows concordance in the individuals affected (aka the mode of inheritance). (Supplemental Figure 5b). Without concordance we do not expect the ASA to be regulating the ASE.

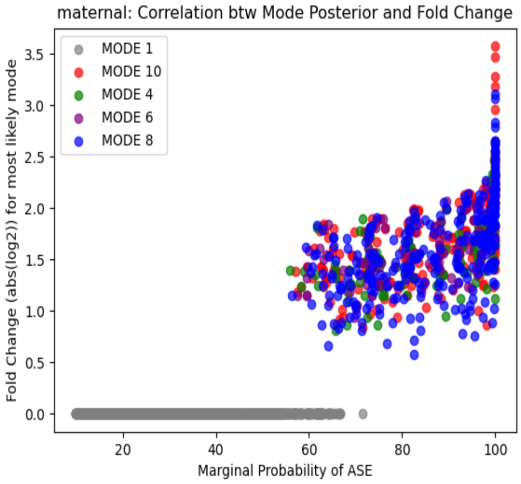


| Gene | Mode | Posterior | Theta | Peak | Mode | Posterior | Theta |
|---|---|---|---|---|---|---|---|
| DDX18 (ENST00000263239) | 10 | 86.78% | 2.69 | chr2:117814454-117815465 (PEAK9784) | 8 [6] [4] | 33.98% [32.51%] [31.68%] | 0.2 [4.71] [6.64] |
| ALDH5A1 (ENST00000357578) | 10 [6] | 51.85% [33.7%] | 0.45 [2.49] | chr6:24522109-24522794 (PEAK16162) | 4 [1] | 48.04% [38.66%] | 0.39 [1] |
| **ACCS (ENST00000263776)** | **8** | **61.66%** | **3.65** | **chr11:44065610-44066611 (PEAK3176)** | **8** | **94.78%** | **2.75** |
| CADM1 (ENST00000331581) | 10 [6] | 62.89% [20.5%] | 3.98 [0.37] | chr11:115194125-115194526 (PEAK3669) | 6 | 79.42% | 4.20 |

**Supplemental Figure 5**: a) ASA peaks identified in the NA12878 trio. B) Peak-gene links in the NA12878 trio. Results are not very concordant, even when looking at secondary modes.
